# Single Serine on TSC2 Exerts Biased Control over mTORC1 Activation by ERK1/2 but Not Akt

**DOI:** 10.1101/2021.07.13.452249

**Authors:** Brittany L. Dunkerly-Eyring, Miguel Pinilla-Vera, Desirae McKoy, Sumita Mishra, Maria I. Grajeda Martinez, Christian U. Oeing, Mark J. Ranek, David A. Kass

## Abstract

The mammalian target of rapamycin complex 1 (mTORC1) is tightly controlled by tuberous sclerosis complex-2 (TSC2) that is regulated by phosphorylation from kinases responding to environmental cues. Protein kinase G specifically modifies serine-1365 (S1364, human), and its phosphorylation (or phosphomimetic SE mutant) potently blocks mTORC1 co-activation by pathological stress, while a phospho-silenced (SA) mutation does the opposite. Neither alter basal mTORC1 activity. Here we show S1365 exerts biased control over mTORC1 activity (S6K phosphorylation) modifying ERK1/2 but not Akt-dependent stimulation. Whereas mTORC1 activation by endothelin-1 is potently modified by S1365 status, insulin or PDGF stimulation are unaltered. TSC2-S1365 is also phosphorylated upon ET-1 but not insulin stimulation in a PKG-dependent manner, revealing intrinsic bias. Neither energy or nutrient modulation of mTORC1 are impacted by S1365. Consistent with these results, knock-in mice with either TSC2 SA or SE mutations develop identical obesity, glucose intolerance, and fatty liver disease from a high fat diet. Thus, S1365 provides an ERK1/2-selective mTORC1 control mechanism and a genetic means to modify pathological versus physiological mTOR stimuli.

## Introduction

The mechanistic target of rapamycin complex 1 (mTORC1) controls cell growth, synthetic activity, metabolism, and protein homeostasis (Liu & Sabatini, 2020; Valvezan & Manning, 2019). Activation of mTOR is tightly regulated by the G-protein Rheb (Ras homolog enriched in brain) which upon GTP binding stimulates mTORC1. This in turn is negatively controlled by the GTPase-activating protein (GAP) tuberin (tuberous sclerosis complex-2, TSC2) (Huang & Manning, 2008). TSC2 complexes with TSC1, and their influence over mTORC1 activity in either direction is prominently regulated by TSC2 phosphorylation from multiple serine/threonine kinases(Huang & Manning, 2008). Prominent kinases that release constitutive TSC inhibition to activate mTORC1 are Akt(Inoki *et al*, 2002; Menon *et al*, 2014) and extracellular response kinase (ERK1/2)(Ma *et al*, 2005). Each modifies specific multiple serines (5 for Akt, 3 for ERK1/2), all but one residing in amino acids 500-1150 of TSC2. TSC2-mediated reduction of Rheb-GTP binding is enhanced by phosphorylation of serines within amino acids 1300-1450 by AMP activated protein kinase (AMPK), glycogen synthase kinase 3β (GSK3β)(Inoki *et al*, 2006), and cGMP-stimulated protein kinase G (PKG)(Ranek *et al*, 2019).

The multitude of different and potentially opposing inputs all convergent on TSC2 raises questions as to how they interact and with what hierarchy. Prior studies have established that a dominant control mechanism involves altered intracellular localization of TSC1/TSC2 at or away from the lysosomal membrane where Rheb and mTORC1 both reside(Carroll *et al*, 2016; Demetriades *et al*, 2016; Fitzian *et al*, 2021; Menon *et al*., 2014). Nutrient starvation or depletion of growth stimuli sends the TSC complex to lysosomes in a TSC1-dependent manner(Fitzian *et al*., 2021) to inhibit mTORC1. By contrast, the complex translocates to the cytosol with Akt stimulation to stimulate mTORC1, though this is blocked with concomitant nutrient depletion(Menon *et al*., 2014). Far less is known regarding how different phosphorylation sites on TSC2 themselves selectively regulate one another, and whether they provide amplitude modulation or more of an on/off switch. An example of the latter is AMPK activation that depresses mTORC1 even in the presence of growth stimuli(Inoki *et al*, 2003). More nuanced control seems likely given system needs to balance multiple signaling inputs.

Proof of a regulatory influence by various kinase modifications of TSC2 were each initially confirmed by genetic mutations to mimic or block phosphorylation. In one or the other direction, these manipulations altered resting mTORC1 activity(Inoki *et al*., 2002; Inoki *et al*., 2006; Ma *et al*., 2005; Manning *et al*, 2002; Potter *et al*, 2002), that made studying their interactions harder to explore. An exception are conserved tandem serines in TSC2 at S1364 or S1365 (human, S1365, 1366 in mouse) phosphorylated by PKG(Ranek *et al*., 2019). If either is mutated to an alanine (SA, phospho-null) or a glutamic acid (SE, phospho-mimetic), basal mTORC1 activity remains unchanged *in vitro* and *in vivo* (Ranek *et al*., 2019). However, upon co-activation by a G_q,11_-GPCR agonist such as endothelin-1 (ET-1), or by pathological pressure stress in intact hearts, mTORC1 activity is greatly amplified if the SA mutation is expressed and equally potently attenuated by the SE mutation (Ranek *et al*., 2019). Here, the major mTORC1 effectors modified pathological muscle growth, hypertrophy, and autophagy, though other studies have revealed control over metabolism in ischemic-reperfusion injury (Oeing *et al*, 2021). Thus, these serine modifications act as rheostats, amplifying or attenuating TSC2-modulated mTORC1 co-activation but not basal activity, and conferring similar potency in either direction. These features could prove useful for evolving cell-therapies where bi-directional mTORC1 modulation and regulation of mTOR that does not risk losing its homeostatic roles as occurs with allosteric and catalytic inhibitors (Pallet & Legendre, 2013) are desired.

In this study, we tested whether TSC2-S1365 broadly modulates mTORC1 co-stimulation or exhibits selectivity in its regulation. Using TSC2 SA and SE mutants, we find that S1365 provides bi-directional modulation of ERK1/2-dependent stimuli yet has negligible influence over Akt-mediated activation of mTORC1. This is paralleled by differences in intrinsic S1365 phosphorylation, and lack of modulation of diet-induced obesity and liver steatosis. S1365 also has negligible influence on mTORC1 activity in response to ischemic conditions, or altered nutrient/energy supply. Thus, TSC2 S1365 provides novel biased bi-directional modulation of mTORC1 activity when co-activated by ERK1/2 but not Akt or nutrient signaling.

## Results

### TSC2 S1365 bidirectionally regulates ERK1/2 activation of S6K

The TSC2 SA mutant amplifies ET-1 stimulated mTORC1 in cardiac myocytes whereas TSC2 SE attenuates the response(Ranek *et al*., 2019). Here we tested the specific role played by ERK1/2 activation. NRMVs were exposed to ET-1 ± a selective ERK1/2 inhibitor (SCH772984, vehicle, or 0.01-10 µM; Figure 1A). ET-1 induced S6K (S389) phosphorylation that corresponded with ERK1/2 but not Akt phosphorylation (the latter at T308 relevant to mTORC1 activity). Inhibiting ERK1/2 blocked the ET-1 S6K response in a dose-dependent manner without altering ERK1/2 phosphorylation. ERK1/2 inhibition also dose-dependently increased Akt T308 phosphorylation likely by a feedback signal, but this had no apparent impact on pS6K that still declined to baseline. By contrast, ET-1 stimulated S6K was not significantly affected by the allosteric pan-Akt inhibitor MK2206 (0.15-15 µM, Figure 1B), leaving ERK1/2 phosphorylation unchanged while reducing Akt T308 phosphorylation(Iida *et al*, 2013).

**Figure 1).**
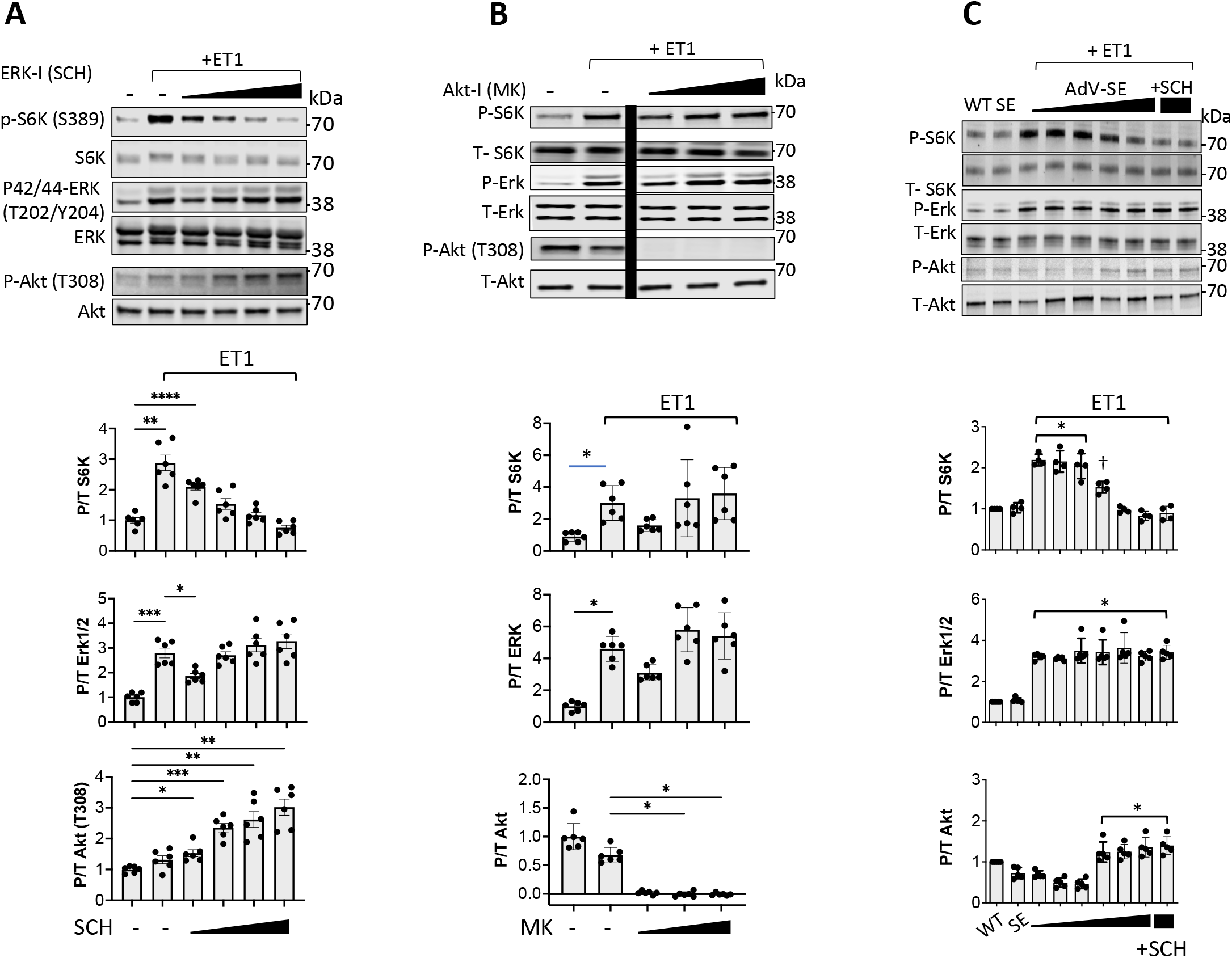
ET-1 stimulates S6K by ERK1/2 Activation blocked by TSC2-S1365E. **A)** Example immunoblots and summary data of dose dependent decline in ET-1 induced S6K phosphorylation by ERK1/2 inhibition (SCH) in NRVMs. ERK1/2 phosphorylation does not significantly change but Akt phosphorylation rises at higher doses. N=6/group; **** P<1e^-4^, *** P≤ 0.0006, ** P≤ .006, * P=0.01; Welch ANOVA, Dunnet’s test for multiple comparisons. **B)** Inhibition of Akt does not significantly reduce S6K activation by ET-1. * p=0.002 Mann Whitney; for pAktp-AktpAkt; N=6/group; Kruskal Wallis test, Dunns MCT; * P = 0.02. **C)** Expression of TSC2-SE mutant reduces ET-1 stimulated S6K in dose dependent manner; ERK1/2 activity is unchanged, and Akt activity rises with higher SE expression. N= 4/group; ANOVA, Sidak’s MCT: * P<10^−7^ versus WT vehicle control; † P= 0.003.

We next tested the dose-dependent impact of TSC2 phospho-state mutations on these responses. ET-1 stimulated S6K declined markedly with incremental SE expression (Figure 1C) as ERK1/2 phosphorylation remained elevated and unchanged, and Akt phosphorylation again increased modestly. By contrast, SA expression resulted in amplified pS6K from ET-1 at the same ERK1/2 activation level, and this was fully blocked by either inhibiting ERK1/2 or mTOR (rapamycin) in cells expressing WT or SA TSC2 (Figure 2A, 2B).

**Figure 2.**
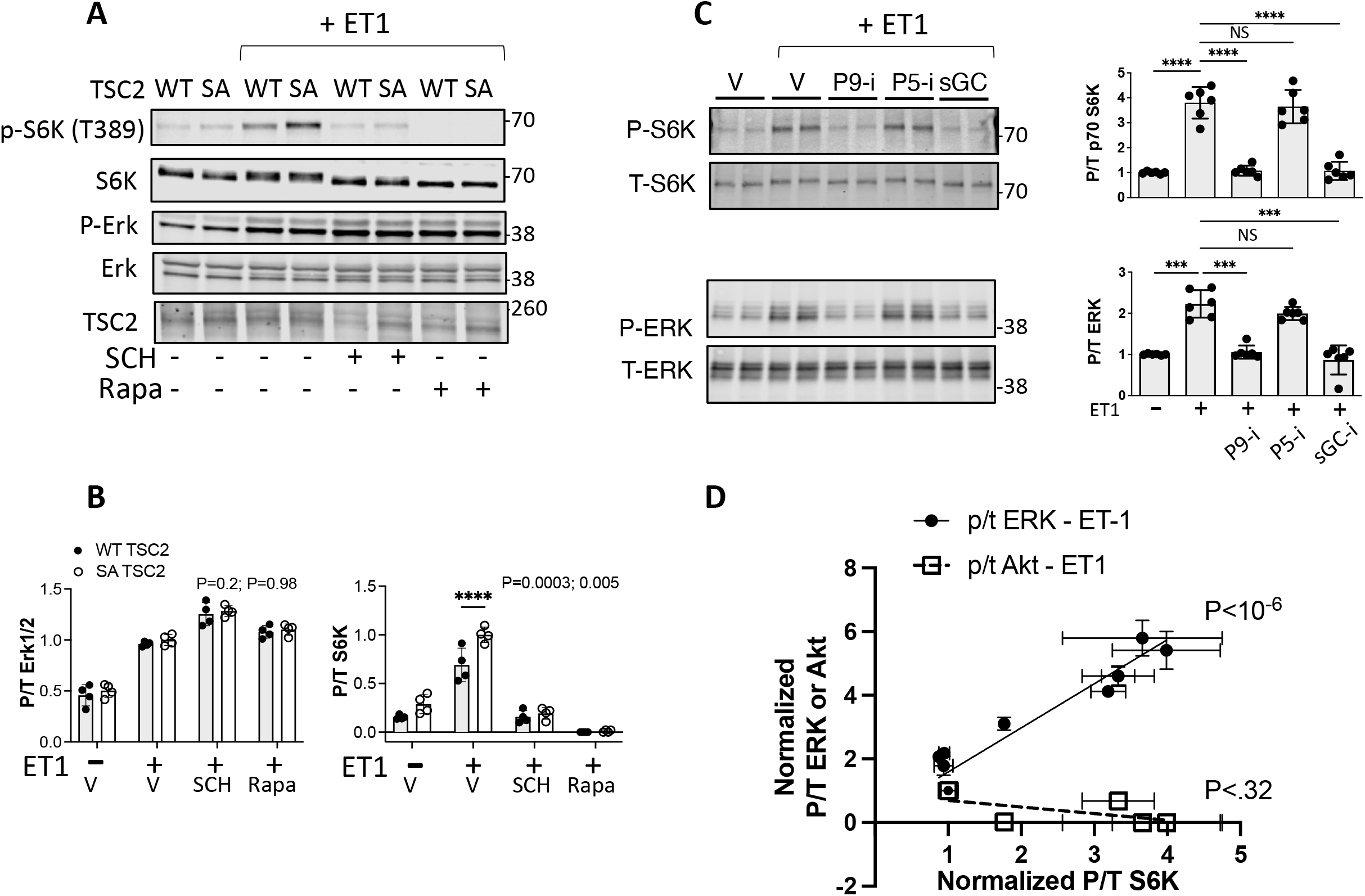
ET-1 stimulated S6K amplified by TSC2-S1365A is ERK1/2 dependent. **A)** Example immunoblots and **B)** summary data showing ET-1 stimulated S6K in NRVMs is augmented by expression of a S1365A (SA) mutation without altering ERK1/2 phosphorylation. This is equally blocked by ERK1/2 (SCH) or mTOR (rapamycin, Rapa) inhibition. N=4/group; P values from 2-way ANOVA for genotype and interaction of genotype and condition. **** p<0.0001 by MCT. **C)** Effect of ET-1 stimulation on S6K and ERK1/2 phosphorylation in NRMVs co-expressing TSC2-S1365A and PKG C42S mutations exposed to PKG activation by different mechanisms. Cells are treated with vehicle (Veh), PDE5 (P5-i) or PDE9 (P9-i) inhibitor, or soluble guanylate cyclase activator (sGC). P9-i and sGC but not P5-i suppress pS6K that corresponds to differential ERK1/2 activation and occurs despite the TSC2-SA mutation that prevents pS1365 modification. N=6/group; Welch ANOVA, Dunnett’s MCT; *** P≤ 0.001; **** P=0.00006. **D)** Summary analysis from data in Figure 1B and 2A revealing how S6K phosphorylation correlates with ERK1/2 but not Akt phosphorylation. Data are replot with baseline normalized to 1.0 and peak ET-1 response to 4.0. P values displayed are for linear regression of each respective relation.

To further explore the importance of ERK1/2 activation, we examined a setting where ET-1 stimulated S6K was differentially suppressed by co-activation of PKG. We compared inhibitors of cGMP-selective phosphodiesterase type 5 (PDE5A), type 9 (PDE9A), or direct soluble guanylate cyclase stimulators, each activating PKG. We recently showed the latter two but not PDE5A inhibition suppress pS6K activation by ET-1 even when the TSC2 SA mutation is present (Oeing *et al*, 2020). This was initially surprising, as PKG is activated by each method, but they do so in different intracellular domains(Lee *et al*, 2015; Nakamura *et al*, 2018); PDE5A is cytosolic whereas PDE9A and sGC also localize to the sarcolemmal membrane(Lee *et al*., 2015; Nakamura *et al*., 2018; Tsai *et al*, 2012) where the ET-1 receptor resides. We hypothesized this might translate to differentially blockade of ERK1/2 activity proximal to TSC2, that would obviate the role of TSC2 S1365. This was tested in NRVMs expressing the TSC2 SA mutation and a redox-dead PKG (C42S) to enhance localization to the sarcolemmal membrane (Nakamura *et al*, 2015). Cells were stimulated by ET-1 to activate S6K, and this activity was unchanged by PDE5A inhibition yet prevented by PDE9-I or sGC activation. There were matching disparities in ERK1/2 activation, being reduced by the latter interventions but not PDE5A inhibition (Figure 2C). Figure 2D combines results from experiments in Figures 1B and 2A, showing a significant dose-response relation between S6K and ERK1/2 but not Akt activity. Together, these studies identify ERK1/2 is a major S6K co-stimulant bi-directionally regulated by TSC2 S1365 (SE or SA mutation).

### TSC2 S1365 minimally reguates Akt-stimulated S6K

We next tested if physiological growth factors that are biased to activate Akt more than ERK/1/2 are also regulated by S1365. Two tyrosine-kinase growth factor agonists - insulin and PDGF were examined. Insulin activated Akt and S6K in mouse embryonic fibroblasts (MEFs), while also co-activating ERK1/2 (Mohan *et al*, 2017) to a much lesser extent (Figure 3A). Importantly, blocking ERK1/2 did not change pS6K, whereas it returned to baseline levels from Akt inhibition, identifying the latter as the primary stimulus. The insulin response was then examined in TSC2 KO-MEFs infected by adenovirus expressing empty vector, or WT, SE, or SA TSC2. KO-cells had higher basal S6K activation that fell similarly by expression of any of the TSC2 genotypes. Insulin induced S6K activation, yet strikingly, there was no decline in cells expressing SE (Figure 3B). This contrasts to full suppression of insulin-stimulated S6K by either an allosteric (rapamycin) or catalytic (torkinib) mTOR inhibitor (Figure 3C). Cells expressing the SA mutation exhibited a modest increase in pS6K over WT, that could reflect co-activated ERK1/2. In each condition, Akt phosphorylation and level of exogenous TSC2 protein expressed were similar. The insulin results were further confirmed using another tyrosine kinase receptor ligand – platelet derived growth factor (PDGF, 20 ng/mL x 30 min) in the same cell model. PDGF also induced robust pAkt and pS6K and both were unaltered by the TSC2 S1365 genotype co-expressed (Figure 3D). Together, these findings show that mTORC1 activation via Akt-dominant pathways is little modified by S1365 TSC2.

**Figure 3.**
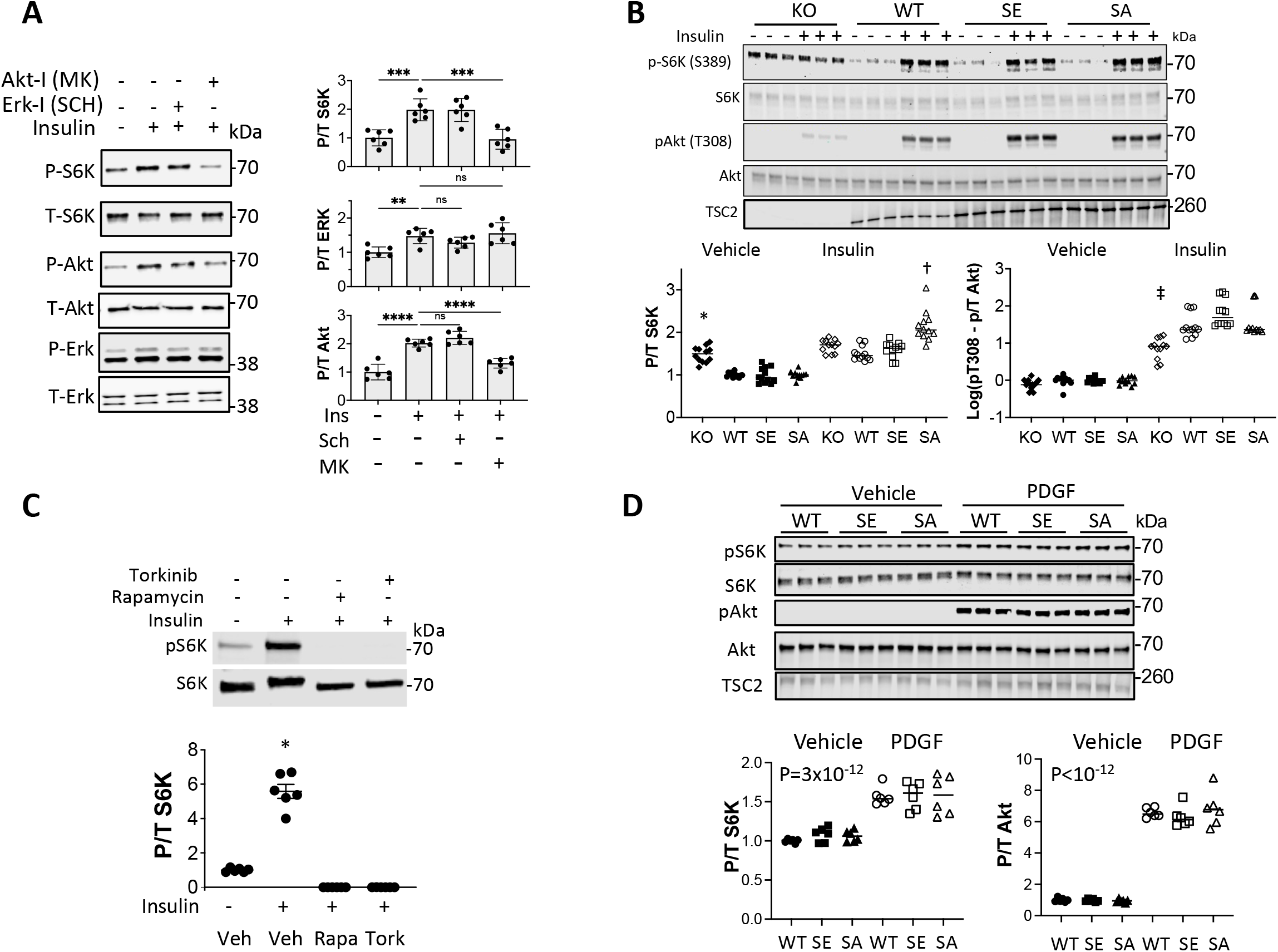
S1365 modulation does not regulate Akt-stimulated pathways. **A)** Insulin stimulated pS6K in MEFs is not altered by ERK1/2 inhibition but is by Akt inhibition; example immunoblot (left) and summary data (right) shown. N=6/group; ANOVA, Holm Sidak MCT: ** P ≤ 0.004, *** P=0.0002; **** P**** P≤ 0.00002. **B)** Example immunoblot and summary results for insulin stimulation in TSC2 KO-MEFs infected with AdV expressing empty vector (KO), or WT, SA, or SE TSC2. N=12/group; 2-Way ANOVA, Sidak’s MCT; * P<2^-7^ versus Vehicle KO; † P<3^-7^ vs KO+ insulin; ‡ P≤6^-7^ versus KO+insulin. **C)** Insulin stimulated S6K is fully blocked by rapamycin (Rapa) or torknib (Tork), contrasting to effect from SE mutant. N=6/group; Welch ANOVA, Dunnet’s test; * p<0.0003 vs other groups. **D)** PDGF stimulation potently activates Akt and S6K and this response is not significantly altered by TSC2 mutants vs WT. N=6/group; 2WANOVA, P=2.6×10^−12^ for PDGF effect, 0.7 for genotype effect, and 0.8 for genotype-PDGF interaction.

### Influence of S1365 modulation on balanced ERK1/2 and Akt co-stimulation

To further assess biased TSC2 S1365 regulation of ERK1/2 versus Akt, we examined thrombin which co-activates both pathways similarly in MEFs (Figure 4A). Cells lacking TSC2 had no S6K activation from thrombin despite ERK1/2 activation, but pS6K rose in cells re-expressing WT-TSC2 (Figure 4B). Blocking Akt reduced thrombin-stimulated pS6K in WT but not the KO cells, while pERK1/2 was unchanged. Alternatively, blocking ERK1/2 in TSC2-WT expressing cells (Figure 4C) reduced thrombin-stimulated pS6K without changing pAkt, supporting dual contributions from both pathways. We then tested if TSC2 SE expression that potently blocks ERK1/2 stimulated S6K also altered Akt activity if both were similarly co-stimulated. While SE expressio reduced the rise in pS6K versus WT (Figure 4D, 4E), reduction in pS6K from Akt inhibition was achieved similarly in each group (Figure 4F). This further supports a negligible impact of S1365 on Akt signaling when co-activated with ERK1/2.

**Figure 4.**
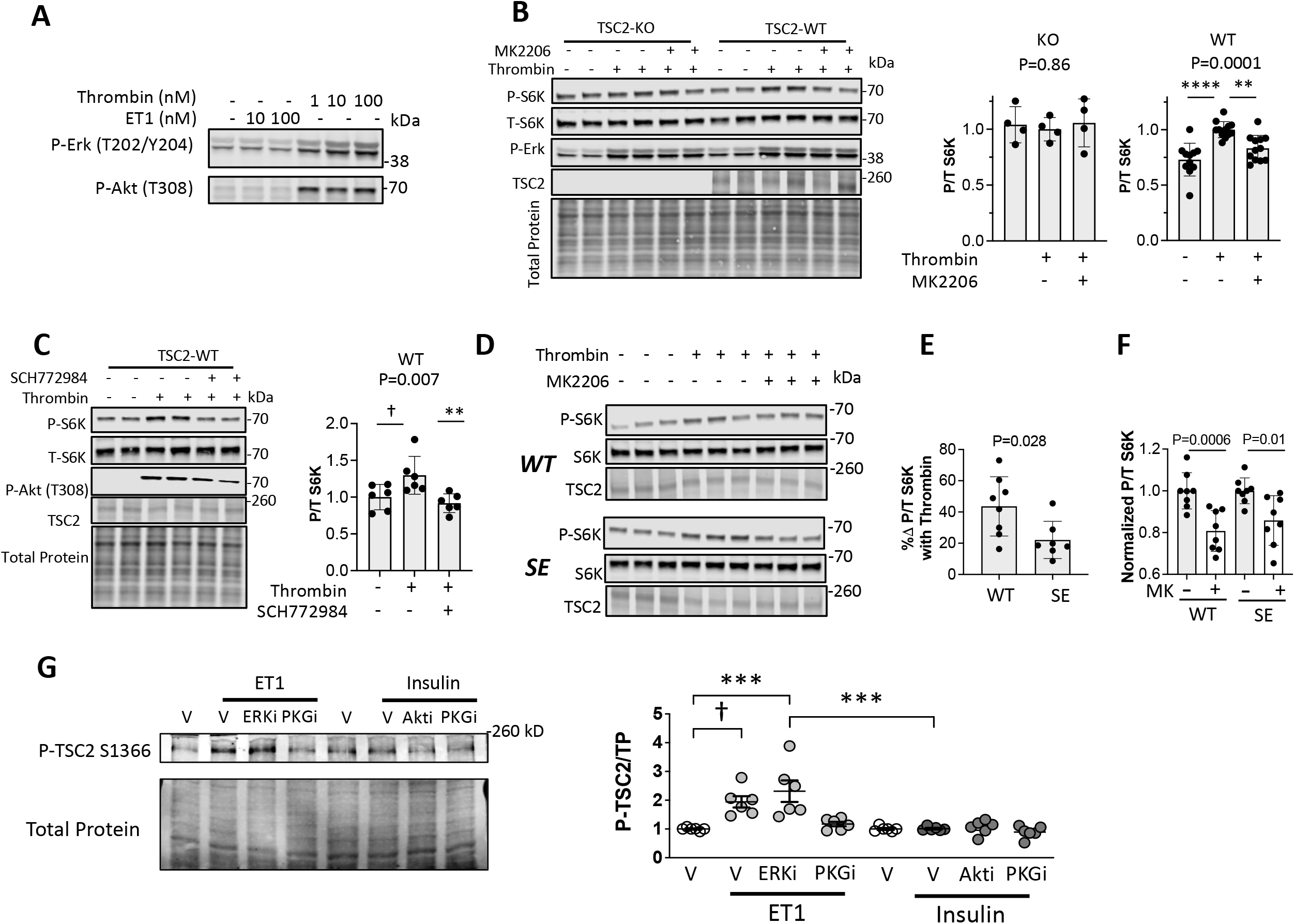
Effect of S1365 on ERK1/2 - Akt Co-activation and impact of each on pS1365. **A) I**mmunoblot of MEFs exposed to thrombin showing balanced activation of ERK1/2 and Akt. **B) I**mmunoblot and summary data for TSC2 KO-MEFs and cells with WT TSC2 re-expressed, exposed to thrombin ± Akt inhibition (MK2206). N=4/group TSC2-KO; N=11/group WT. P values displayed are for Kruskal Wallis test, **** P= .00009, ** P=0.009 by Dunns’ MCT. **C)** Example immunoblot and summary data for the same experiment but ± ERK1/2 inhibitor (SCH). N=6/group; KW test P-values displayed; Dunns MCT: † P=0.1, ** P=0.007. **D)** Example immunoblot for KO-MEFs re-expressing either TSC2-WT or TSC2-SE, stimulated with thrombin ± Akt inhibitor (MK2206). **E)** Summary data from this experiment showing percent rise in P/T S6K from thrombin in SE vs WT TSC2 expressing KO-MEFs. N=7-8/group; P value displayed Mann Whitney U test. **F)** Effect of Akt inhibition on P/T S6K in response to thrombin in cells expressing WT versus SE TSC2. N=8/group; 2W-ANOVA, Sidak’s MCT - P-values displayed. **G)** Differential effect of ET1 or insulin stimulation on TSC2 pS1366 (equivalent to S1365 in mouse), and impact of ERK1/2 or Akt inhibition, or PKG activation. N=6/group; 1WANOVA, Sidak’s MCT. * P=0.004; ** P=0.002; † - p=0.0027; ‡ P=0.018; § P=0.94.

### Increased pS1365 is coupled to ET-1 but not insulin stimulated mTORC1

The amplification by the SA mutation of mTORC1 activation by ET1-ERK1/2 but not insulin-Akt stimulation suggested pS1365 is normally engaged as a counter-brake by the former but not latter pathway. To test this, we studied NRVMs exposed to either ligand for 15 min, with vehicle, ERK1/2 or Akt inhibitor respectively, or the PKG inhibitor DT3 (1 µM) which is known to phosphorylate the site in cardiomyoctes(Ranek *et al*., 2019). S1365 phosphorylation increased after ET-1 stimulation, and this was unaltered with ERK1/2 inhibition but prevented by PKG inhibition (Figure 4G). By contrast, Ins did not increase pS1365, nor was this modified further by adding either Akt or PKG inhibitors. Thus ET-1 but not Ins stimulation results in PKG-dependent S1365 phosphorylation, supporting intrinsic differential regulation of the pathways.

### S1365 minimally controls mTORC1 via energy/nutrient depletion/repletion

Energy depletion activates AMPK, and its phosphorylation of TSC2 at S1387 suppresses S6K activity equally in TSC2^WT^ or TSC2^SA^ expressing KO-MEFs (Ranek *et al*., 2019). Here we tested a simulated ischemia (SI) model combining reduced glucose, acidosis, and hypoxia that stimulates AMPK (Mungai *et al*, 2011) and REDD1 (protein regulated in development and DNA damage) (DeYoung *et al*, 2008) to reduce S6K activation. Figure 5A shows marked reduction in S6K phosphorylation with SI that was negligibly altered by expressing TSC2 SA or SE.

**Figure 5.**
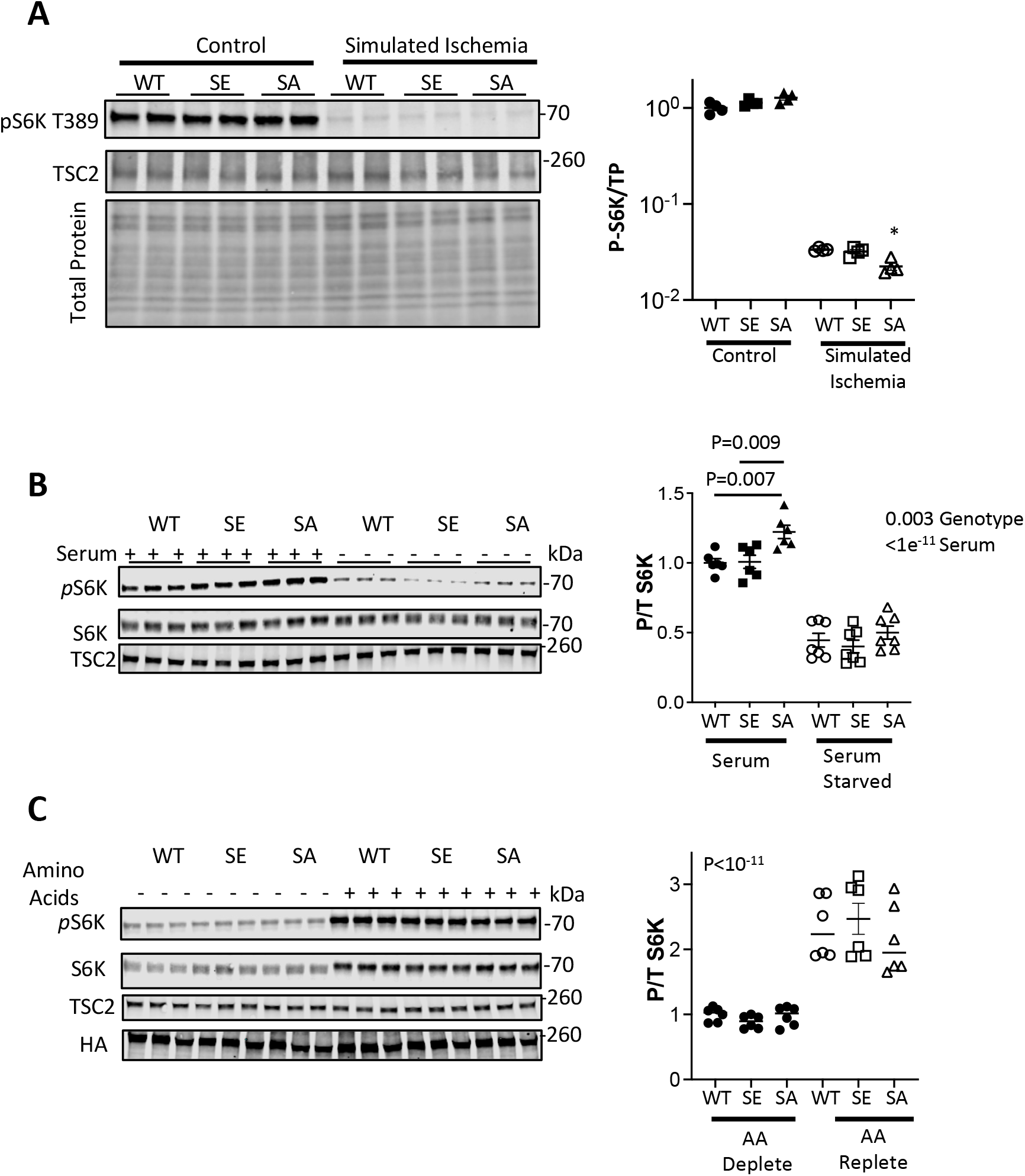
TSC2 S1365 has minimal influence on energy or nutrient regulation of S6K. **A)** NRMVs exposed to simulate ischemia (SI) display marked suppression of S6K activation that is similar regardless of the TSC2 S1365 genotype. There is slightly less activation after SI in SA expressing cells. N=4/group; Kruskal Wallis test, Dunns MCT: * P=0.03 vs WT-SI. **B)** Effect of serum depletion on S6K activation. With serum, pS6K was somewhat greater in SA expressing cells, but all decline markedly with serum depletion regardless of genotype. N=6-7/group; 2WANOVA P-values provided; Sidak’s MCT: P values shown. **C)** Amino acid repletion stimulates S6K similarly regardless of S1365 genotype expressed. P value for 2WANOVA genotype effect displayed. Effect of genotype: P= 0.72; Genotype x AA repletion - P=0.44.

Depletion or abundance of growth factors and/or nutrients are also potent modulators of mTORC1 activity by altering TSC2 localization to or from the lysosome, respectively (Demetriades *et al*., 2016; Menon *et al*., 2014). To test the impact of TSC2 S1365 modification to these stimuli, NRVMs expressing each form were cultured in FBS followed by serum-free conditions (each 1 hr). Cells in FBS had slightly higher S6K activity if they expressed SA (Fig. 5B) consistent with the presence of multiple growth factors. However, with serum depletion, levels declined markedly and independent of TSC2 genotype. We also tested the effects of amino acid depletion, with cells placed in dialyzed FBS removing free amino acids, and then switched to media containing amino acids (each 1 hr). S6K activation was low with amino acid depletion and increased similar in all TSC2 groups with its repletion (Figure 5C). Together, these data indicate that TSC2 1365 modification does not influence energy or nutrient stimuli associated with S6K activation or inhibition.

### S1365 phosphosite mutagenesis does not alter diet-induced obesity

In hearts subjected to either pressure-overload and ischemia-reperfusion injury, the status of S1365 (e.g. SA or SE) potently alters the consequent levels of mTORC1 activity and corresponding pathophysiology(Oeing *et al*., 2021; Ranek *et al*., 2019). Both stresses engage ERK1/2 stimulation, the former in conjunction with activation of multiple pathogenic kinases(Bueno *et al*, 2000), and the latter as a protective mechanism(Lips *et al*, 2004), both directions being concordant with the detriment or benefit respectively from SA expression. Another stress, diet-induced obesity, also stimulates mTORC1 in fat, liver, and skeletal muscle(Khamzina *et al*, 2005); however, this is primarily achieved by Akt(Dann *et al*, 2007; Khamzina *et al*., 2005; Zhang *et al*, 2009b) - S6K signaling to stimulate adipogenesis and obesity, fatty liver, and metabolic syndrome(Jia *et al*, 2014; Um *et al*, 2004). Based on the current results, we hypothesized that mice harboring a WT, SA or SE TSC2 mutation would respond similarly to diet-induced obesity.

After 18 weeks of a HFD, each genotype developed superimposable increases in body weight (Figure 6A). Resting plasma glucose was also similarly elevated in each group (P=and the response to a glucose challenge was essentially identical (Figure 6B). In all models, there was extensive lipid deposition in the liver in marked contrast to livers from aged-matched mice in each group fed a control diet (Figure 6C). Together, these results in a relevant model known to involve Akt-mediated mTORC1 activation as a major component of its pathophysiology was unaltered by SA or SE mutations.

**Figure 6.**
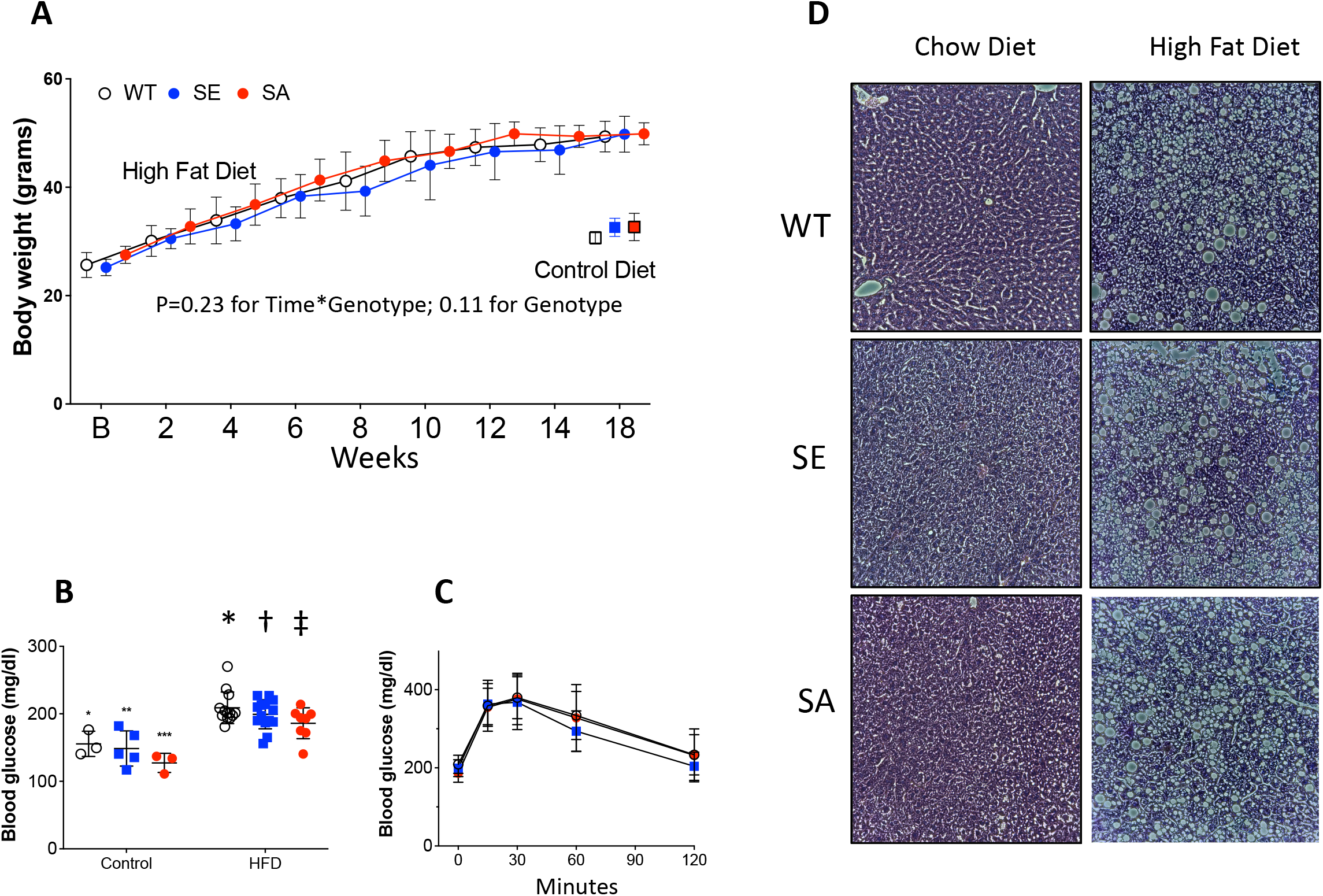
TSC2 S1365 does not regulate response to diet induced obesity. **A)** Body weight increased similarly in SA or SE homozygous KI mice or littermate controls over 18 week duration. N: Control: WT=3, SE=5, SA-3, high fat diet (HFD): WT=14; SE=14, SA=8; 2WANOVA, P=0.23 for time x genotype; p=0.11 for genotype; P<10^−10^ for time. **B)** Basal glucose is similarly elevated in each genotype group. Replicates as in panel A. ANOVA: P=0.9 for genotype x diet interaction; HFD vs control: * P=0.002, † P=0.0003; ‡ P=001. **C)** Glucose response test also displayed very similar responses indicating similar insulin desensitization. N=6/group; 2WANOVA: P=0.31 for time x genotype, 0.57 genotype, P=10^−9^ time). **D)** Examples of H/E-stained liver from WT, SA, and SE KI mice with control versus HFD. There is similar marked steatosis independent of the TSC2 genotype. Controls on standard diet are shown below each panel. Data reflect total n=6-8/group, with P>0.6 for genotype effect on steatosis (χ^2^ test).

## Discussion

Modulation of mTORC1 activation by TSC2 involves balancing multiple signaling inputs, the majority being transduced by phosphorylation at specific serines and threonines within a 900 amino acid mid-region of the protein. Unlike Akt, ERK1/2, AMPK, or GSK3β that require altering multiple residues to modulate mTORC1 (Inoki *et al*., 2002; Inoki *et al*., 2006; Inoki *et al*., 2003; Ma *et al*., 2005; Manning *et al*., 2002), modification of one serine at S1365 is sufficient to confer regulatory effects – not on basal but on co-stimulated mTORC1 activity. Here we show this regulation applies to ERK1/2 but not Akt activation pathways, nor does it modulate mTORC1 modulation by nutrient or amino-acid supply. The latter differs from Akt-TSC2 phosphorylation and S6K activation(Menon *et al*., 2014) that is blocked by nutrient depletion. Figure 7 summarizes the signaling revealed in our study. While we tested several growth factors, there are many receptor tyrosine kinases, cytokines, immune antigen receptors, and G-protein coupled receptors that also stimulate Akt(Manning & Toker, 2017) We suspect these are also unlikely to be altered by S1365 modulation. Alternatively, many pathways that engage ERK1/2, including GPCRs, oxidant stress, protein kinase C and RSK1 stimulation (Ma *et al*., 2005) are likely modified by S1365. These findings reveal a new means of nuanced mTOR1 regulation via TSC2 to regulate some forms of co-activation but not others.

**Figure 7.**
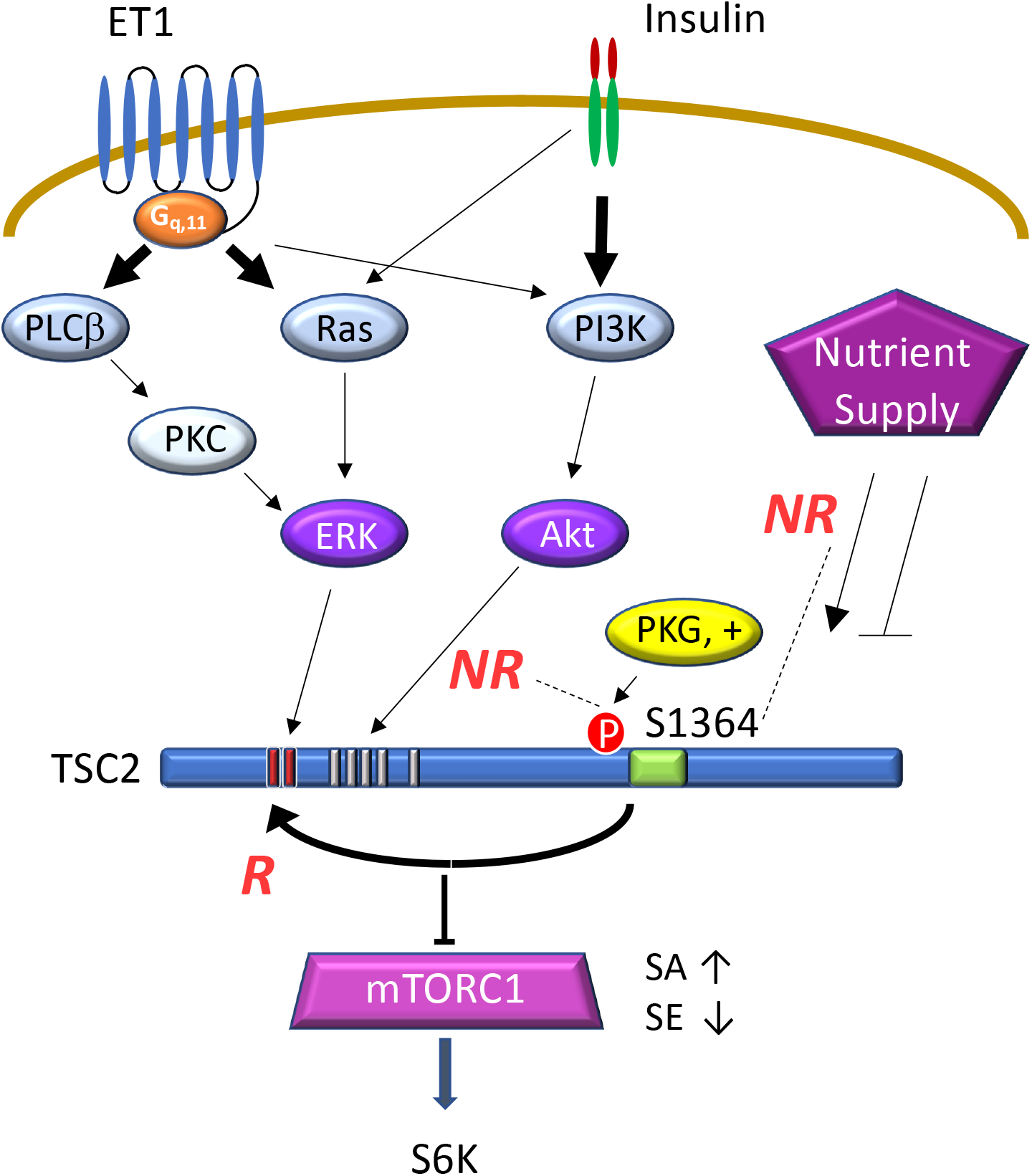
Schematic of TSC2-1365 regulated and non-regulated signaling. A G_q,11_ – GPCR typified by ET-1 receptor and TKR typified by insulin receptor are depicted along with their downstream signaling. ET-1 prominently activated PLCβ-PKC-ERK and Ras-ERK pathways leading to TSC2 phosphorylation and mTORC1 activation indexed by pS6K. Insulin more prominently activates PI3K-Akt also leading to mTORC1 activation. However, the former but not latter results in concomitant phosphorylation of TSC2 S1365. An SA mutation of this site blocks this and amplifies ET-1 but not insulin mediated mTORC1 activation, while an SE mutation mimics pS1365, also blunting ET-1 but not insulin stimulated mTORC1. ET-1 stimulated pS1365 is PKG but not ERK-dependent and involves kinase activation upstream of ERK as it is unaltered despite ERK1/2 inhibition. This correlates with the capacity of S1365 modulation to potently alter ET-1 activation of mTORC1. By contrast, insulin does not lead to pS1365 and its activation of mTORC1 is correspondingly insensitive to S1365 phospho-mutations.

The strong bias for S1365 to control ERK1/2 but not Akt modulation of mTORC1 is consistent with its intrinsic phosphorylation by the former but not latter stimulus. Gq-coupled GPCRs activate ERK1/2 by phospholipase Cβ-PKC and Ras dependent pathways. Both ET-1(Kapakos *et al*, 2010; Nakamura *et al*., 2015) and Ins(Anfossi *et al*, 2009; Yu *et al*, 2011) co-activate nitric oxide synthase dependent cGMP-PKG activation via Akt-dependent signaling. Yet we only observed pS1365 with ET-1 stimulation. This is intriguing and may result from locally constrained PKG signaling, a property of the kinase. It indicates S1365 is likely an intrinsic and selective counter-brake on certain mTORC1 activation pathways, with those that themselves lead to enhanced pS1365 being modulated by its phospho-status (or site mutagenesis) and those that do not alter pS1365 little impacted. PKG is not the only kinase that can phosphorylate S1365, as PKC can do this as well(Ballif *et al*, 2005), and other kinases may also play a role in other cell types.

Evidence to date shows that rather than altering TSC2-GAP activity, the primary mechanism by which TSC2 modulates mTORC1 is by altering intracellular localization to or away from the lysosomal membrane. As first reported by Manning and Demetriades, starvation and growth factor or amino acid depletion moves TSC2 to the lysosome to reduce Rheb-GTP binding and suppress mTORC1, whereas Akt activation does the opposite(Demetriades *et al*, 2014; Demetriades *et al*., 2016; Menon *et al*., 2014). In 2021, Fitzian et al. reported that TSC1 is important for this translocation, binding to phosphoinositol phospholipid PI3,5P2 at the lysosomal membrane and thereby positioning TSC2 for mTORC1 inhibition. Whether this or other mechanisms apply to TSC complex movement from lysosomes upon Akt or ERK1/2 activation or their selective modulation by S1365 is the subject of ongoing research. However, the lack of basal phenotypes with both SA and SE mutants, and preservation of nutrient, energy, and physiological growth factor signaling via Akt suggests something else is involved. To our knowledge, evidence that ERK1/2 activation translocates TSC1/TSC2 complex to the cytosol is not reported, but if it did, then the SE mutation would appear to reverse it and SA augment the translocation. Mechanisms for such bi-directional and biased control have not yet been identified.

MTORC1 activation in diet-induced obesity (DIO) has long been thought to play a major pathophysiological role, contributing to abnormal tissue fat deposition particularly in liver (Han & Wang, 2018; Wang *et al*, 2015), type-2 diabetes and insulin desensitization(Bar-Tana, 2020), lypogenesis(Zhang *et al*, 2009a), and other changes. Mice lacking S6K are resistant to this stress and consequent metabolic syndrome(Um *et al*., 2004). Yet, none of these abnormalities were altered by preventing or mimicking S1365 phosphorylation. When we first performed this in vivo study, we expected SA mice to exhibit worse DIO and metabolic syndrome, and SE mice the opposite. While the results seemed puzzling at first, they later made sense in light of the dichotomy between S1365 regulation of ERK1/2 but not Akt stimulated mTORC1. They also help clarify potential targets for S1365 modulation, suppressing Gq-GPCR, ROS, and other pathological stimuli, while preserving physiological signaling mediated by insulin and other receptor tyrosine kinases (RTKs). It is interesting that PKG stimulation is known to counter pathological vascular and cardiac growth/remodeling as with chronic hypertension, but not to impair physiological growth adaptations as with exercise(Takimoto *et al*, 2009). Furthermore, though activating PKG suppresses pressure-overload cardiac disease, it does not block analogous pathology due to overexpressed Akt activation(Takimoto *et al*, 2005).

Most of the experiments presented in this study stimulated pathways using receptor ligands as opposed to over-expressing a constitutively active kinase. We chose this approach to generate acute changes rather than prolonged signaling that can itself trigger compensations. While these ligands did not purely trigger one pathway, (ERK1/2 vs Akt), their balance was sufficiently different to address the hypothesis. We also used selective kinase inhibitors rather than genetic silencing methods, as the latter can also induce feedback loops via other signaling to make interpretations less direct. We have not yet identified the intra-molecular mechanism by which S1365 on TSC2 can selectively modulate ERK1/2 but not Akt signaling. However, recent molecular structure data and ongoing efforts should pave the way for such insights in the future. The finding of parallel disparities in the phosphorylation of S1365 with ET-1 but not Insulin stimulation may help identify this mechanism. PKG is a primary S1365-modifying kinase in both cardiomyocytes and fibroblasts that results in suppressing mTORC1 co-activation. Other kinases reported to modify the site such as PKC also activate mTORC1 so this maybe another example of a co-stimulated brake. Control by S1365 of several other TSC2-regulatory pathways, including p38 MAP kinase and 14-3-3 (DeYoung *et al*., 2008; Huang & Manning, 2008) that inhibit TSC2 to activate mTORC1, remains to be determined.

To our knowledge, S1365 is the first TSC2 regulatory site revealed that not only leaves basal mTORC1 signaling intact despite mutagenesis to phospho-silenced or mimetic forms but is also selective to modulating mTORC1 activity when pathological ERK1/2 co-stimulation is present yet preserving signaling coupled to many other modulators. While first revealed in cardiomyocytes and MEFs, this pathway is ubiquitous and likely has regulatory significance in all cells. Only single gene mutations are needed to generate SA or SE forms, making gene editing for adoptive cell therapy quite tractable. Ongoing studies are assessing the use of these gene modifications to enhance immunological treatment of diseases such as cancer, and the present findings help clarify what type of signal amplification/attenuation one can expect.

## Materials and methods

### Cell culture models

Rat neonatal cardiomyocytes (NRVMs) were freshly isolated as previously described(Lee *et al*., 2015) and cultured at 1 million cells per well in six-well plates for 24 hr in DMEM with 10% FBS and 1% penicillin/streptomycin before study. In addition, studies were performed using TSC2 knockout mouse embryonic fibroblasts (MEFs) were cultured in DMEM with 10% FBS and 1% pen/strep to 50-60% confluence. Both NRVMs and TSC2 KO MEFs were modified to express WT, SA, or SE TSC2 mutant protein using either adenovirus or plasmid expression vectors applied at reported MOI(Oeing *et al*., 2020) for 3 to 4 hours in serum free media. Plasmids were applied 24 hours after plating, (5 μg with Takara Clontech Xfect in NRVMs, 2.5 ug and 7.5ul Lipofectamine LTX reagent, Thermo Fisher’s Lipofectamine LTX & PLUS Reagent, MEFs) following manufacturer’s protocol. All cells were provided 48 hours to express the TSC2 proteins prior to study. To assess intrinsic TSC2 S1365 phosphorylation, NRCMs were cultured in no-serum media for 24 hours prior to study, then pre-treated with SCH772984 (10 uM), Akt (MK-2206, 150 nM), PKG (DT3, 1 uM), or vehicle for one hour prior to ET-1 (100 nM), insulin (10 ug/ml), or vehicle stimulation for 15 minutes. They were then lysed and examined for TSC2 S1365 phosphorylation.

### Stimulation and Kinase Modulation

ERK1/2 stimulation was achieved using either endothelin-1 (ET-1) in s, or thrombin in MEFs. Selective ERK1/2 inhibition was achieved with 10 μM SCH772984 and Akt inhibition using 150 nM MK-2206 (unless otherwise indicated), each provided 1 hour prior to stimulation by 100 nM ET-1 or 100 nM thrombin for 15 min. In additional NRVM studies, PKG was stimulated using a selective inhibitor of either phosphodiesterase type 9 (PF-04449613, 0.1 μM), type 5 (sildenafil, 1 μM), or guanylate cyclase activator (Bay-602770, 0.1 μM).

Effects from Akt stimulation was tested using insulin or platelet derived growth factor (PDGF) in MEFS or TSC2-KO MEFs, the latter also infected with AdV expressing WT, SE, or SA TSC2. Cells were serum starved overnight and then stimulated with insulin (0.5 μM) or PDGF (20 ng/ml) for 30 minutes. Doses were based on screening dose response assays assessing for Akt and S6K activation.

Simulated ischemia, serum withdrawal, and amino acid depletion/repletion were studied in NRVMs. To simulate ischemia, NRVMs were subjected to 30 minutes of hypoxia, acidosis, and reduced glucose using a hypoxia chamber and ischemia-mimicking buffer containing 125 mM NaCl, 6.25 mM NaHCO_3_, 1.2 mM KH_2_PO_4_, 1.2 mM CaCl_2_, 1.25 mM MgSO_4_, 5 mM sodium lactate, 8 mM KCl, 20 mM HEPES, and 20 mM 2-deoxyglucose (pH adjusted to 6.6) as described(Oeing *et al*., 2021). For serum withdrawal, cells were provided fresh DMEM supplemented with 10% FBS for 1 hour followed by incubation with DMEM lacking FBS for 1 hour. Amino acid depletion/repletion used DMEM deplete of amino acids and supplemented with 10% dialyzed FBS and 1% pen/strep for 1 hour, followed by medium containing the full complement of amino acids, DMEM, and supplemented with 10% dialyzed FBS and 1% pen/strep.

### Reagents, constructs, and antibodies

Primary antibodies used were anti-phospho-p70 S6 kinase (Thr389) (Cell Signaling Technology, 1:1000), anti-p70 S6 kinase (Cell Signaling Technology, 1:1000), anti-phospho-4E-BP1 (Ser65) (Cell Signaling Technology, 1:1000), anti-phospho-ULK1 (Ser757) (Cell Signaling Technology, 1:1000), anti-4E-BP1 (Cell Signaling Technology, 1:1000), anti-phospho-Akt (Ser473) (Cell Signaling Technology, 1:1000), anti-phospho-Akt (Thr308) (Cell Signaling Technology, 1:1000), anti-Akt (Cell Signaling Technology, 1:1000), anti-Tuberin/TSC2 (Cell Signaling Technology, 1:1000), anti-phospho-p44/42 MAPK (ERK1/2) (Thr202/Tyr204) (Cell Signaling Technology, 1:1000), and anti-p44/42 MAPK (ERK1/2) (Cell Signaling Technology, 1:1000). Total protein stain used was Revert Total Protein Stain (Licor). We generated N-terminally HA tagged TSC2 constructs as previously described (Ranek *et al*., 2019). Adenovirus was developed expressing the HA-tagged wild type human TSC2, TSC2 (S1364A) or TSC2 (S1364E) as previously described (Ranek *et al*., 2019).

### Immunoblotting

Cells were lysed in lysis buffer (Cell Signaling Technology, 9803) and the protein concentrations were determined by BCA assay (Pierce). Samples were prepared in Licor protein sample loading buffer and run on TGX 7.5% and 4-20% Tris-glycine gels (Bio-Rad) and blotted onto a nitrocellulose membrane. Membranes were incubated with primary antibodies overnight at 4°C, followed by incubation with secondary antibodies (Licor). Membranes were imaged with infrared imaging (Odyssey, Licor) and quantified with Licor Image Studio Software 3.1.

### In Vivo High-fat diet induced Obesity

Age matched male mice harboring homozygous KI mutations for SA (n=8), SE (n=14) or littermate WT controls (n=14) were fed 60% high fat diet (Research Diets, D12492, 60 kcal% fat) for 18 weeks, starting at age 10-12 weeks. An additional group of n=3-5 mice of each genotype were placed on regular diet for the same time. Serial weights were obtained every other week. After 18 weeks resting plasma glucose and glucose tolerance tests were obtained after overnight fasting (18 hours). For the glucose tolerance test, the mice injected with 10 μl/gm body weight sterile 10% glucose in normal saline. Blood glucose was then measured serially by standard blood glucose meter from a tail-tip clip-bleed.

### Statistical Analysis

Data are mostly analyzed by 1- or 2-way ANOVA, or by non-parametric Kruskal Wallis or Mann Whitney U tests, as appropriate. Each test is identified in the figure legends. Multiple comparisons tests (Sidak, Dunns, or Tukey) were used to obtain within group comparisons, and area also identified in the legends. Statisitcal analysis was performed using Prism Version 9.0.

## Acknowledgements

This study was supported by National Institute of Health – Heart Lung and Blood Institute grants: (HL135827 (DAK) and HL 143905 (BDE); T32-HL-7227 (MPV); American Heart Association 16SFRN28620000 (DAK) and 18CDA34110140 (MJR); and Deutsche Forschungsgemeinschaft (German Research Foundation) OE 688/1-1 (CUO), BIH-Charité clinical scientist program funded by the Charité – Universitätsmedizin Berlin and the Berlin Institute of Health (CUO).

## Author contributions

BDE performed most of the experiments, analyzed the data and constructed the manuscript. MPV performed the chronic high fat diet study, SM, JP, DM, MIG, CUO, and MJR each provided results for individual cell-based experiments and analyses. DAK designed the studies, performed data analysis and presentation, and edited the manuscript.

## Conflict of interest

DAK, BDE, and MJR are co-inventors on a PCT patent regarding the use of TSC2 S1364/65 (human) mutations for immunological therapeutics.

## Notes

### Competing Interest Statement

David A. Kass, Mark Ranek, and Brittany Dunkerly-Eyring are all co-inventors on filed patents regarding the use of TSC2 site mutation to modulate immune-therapy.

## References

Anfossi G, Russo I, Doronzo G, Trovati M (2009) Contribution of insulin resistance to vascular dysfunction. Arch Physiol Biochem 115: 199–217

Ballif BA, Roux PP, Gerber SA, MacKeigan JP, Blenis J, Gygi SP (2005) Quantitative phosphorylation profiling of the ERK/p90 ribosomal S6 kinase-signaling cassette and its targets, the tuberous sclerosis tumor suppressors. Proc Natl Acad Sci U S A 102: 667–672

Bar-Tana J (2020) Type 2 diabetes - unmet need, unresolved pathogenesis, mTORC1-centric paradigm. Rev Endocr Metab Disord 21: 613–629

Bueno OF, De Windt LJ, Tymitz KM, Witt SA, Kimball TR, Klevitsky R, Hewett TE, Jones SP, Lefer DJ, Peng CF et al (2000) The MEK1-ERK1/2 signaling pathway promotes compensated cardiac hypertrophy in transgenic mice. EMBO J 19: 6341–6350

Carroll B, Maetzel D, Maddocks OD, Otten G, Ratcliff M, Smith GR, Dunlop EA, Passos JF, Davies OR, Jaenisch R et al (2016) Control of TSC2-Rheb signaling axis by arginine regulates mTORC1 activity. Elife 5

Dann SG, Selvaraj A, Thomas G (2007) mTOR Complex1-S6K1 signaling: at the crossroads of obesity, diabetes and cancer. Trends in molecular medicine 13: 252–259

Demetriades C, Doumpas N, Teleman AA (2014) Regulation of TORC1 in response to amino acid starvation via lysosomal recruitment of TSC2. Cell 156: 786–799

Demetriades C, Plescher M, Teleman AA (2016) Lysosomal recruitment of TSC2 is a universal response to cellular stress. Nature communications 7: 10662

DeYoung MP, Horak P, Sofer A, Sgroi D, Ellisen LW (2008) Hypoxia regulates TSC1/2-mTOR signaling and tumor suppression through REDD1-mediated 14-3-3 shuttling. Genes Dev 22: 239–251

Fitzian K, Bruckner A, Brohee L, Zech R, Antoni C, Kiontke S, Gasper R, Linard Matos AL, Beel S, Wilhelm S et al (2021) TSC1 binding to lysosomal PIPs is required for TSC complex translocation and mTORC1 regulation. Mol Cell

Han J, Wang Y (2018) mTORC1 signaling in hepatic lipid metabolism. Protein & cell 9: 145–151

Huang J, Manning BD (2008) The TSC1-TSC2 complex: a molecular switchboard controlling cell growth. Biochem J 412: 179–190

Iida M, Brand TM, Campbell DA, Starr MM, Luthar N, Traynor AM, Wheeler DL (2013) Targeting AKT with the allosteric AKT inhibitor MK-2206 in non-small cell lung cancer cells with acquired resistance to cetuximab. Cancer Biol Ther 14: 481–491

Inoki K, Li Y, Zhu T, Wu J, Guan KL (2002) TSC2 is phosphorylated and inhibited by Akt and suppresses mTOR signalling. Nat Cell Biol 4: 648–657

Inoki K, Ouyang H, Zhu T, Lindvall C, Wang Y, Zhang X, Yang Q, Bennett C, Harada Y, Stankunas K et al (2006) TSC2 integrates Wnt and energy signals via a coordinated phosphorylation by AMPK and GSK3 to regulate cell growth. Cell 126: 955–968

Inoki K, Zhu T, Guan KL (2003) TSC2 mediates cellular energy response to control cell growth and survival. Cell 115: 577–590

Jia G, Aroor AR, Martinez-Lemus LA, Sowers JR (2014) Overnutrition, mTOR signaling, and cardiovascular diseases. American journal of physiology Regulatory, integrative and comparative physiology 307: R1198–1206

Kapakos G, Bouallegue A, Daou GB, Srivastava AK (2010) Modulatory Role of Nitric Oxide/cGMP System in Endothelin-1-Induced Signaling Responses in Vascular Smooth Muscle Cells. Current cardiology reviews 6: 247–254

Khamzina L, Veilleux A, Bergeron S, Marette A (2005) Increased activation of the mammalian target of rapamycin pathway in liver and skeletal muscle of obese rats: possible involvement in obesity-linked insulin resistance. Endocrinology 146: 1473–1481

Lee DI, Zhu G, Sasaki T, Cho GS, Hamdani N, Holewinski R, Jo SH, Danner T, Zhang M, Rainer PP et al (2015) Phosphodiesterase 9A controls nitric-oxide-independent cGMP and hypertrophic heart disease. Nature 519: 472–476

Lips DJ, Bueno OF, Wilkins BJ, Purcell NH, Kaiser RA, Lorenz JN, Voisin L, Saba-El-Leil MK, Meloche S, Pouysségur J et al (2004) MEK1-ERK2 signaling pathway protects myocardium from ischemic injury in vivo. Circulation 109: 1938–1941

Liu GY, Sabatini DM (2020) mTOR at the nexus of nutrition, growth, ageing and disease. Nat Rev Mol Cell Biol 21: 183–203

Ma L, Chen Z, Erdjument-Bromage H, Tempst P, Pandolfi PP (2005) Phosphorylation and functional inactivation of TSC2 by Erk implications for tuberous sclerosis and cancer pathogenesis. Cell 121: 179–193

Manning BD, Tee AR, Logsdon MN, Blenis J, Cantley LC (2002) Identification of the tuberous sclerosis complex-2 tumor suppressor gene product tuberin as a target of the phosphoinositide 3-kinase/akt pathway. Mol Cell 10: 151–162

Manning BD, Toker A (2017) AKT/PKB Signaling: Navigating the Network. Cell 169: 381–405

Menon S, Dibble CC, Talbott G, Hoxhaj G, Valvezan AJ, Takahashi H, Cantley LC, Manning BD (2014) Spatial control of the TSC complex integrates insulin and nutrient regulation of mTORC1 at the lysosome. Cell 156: 771–785

Mohan ML, Chatterjee A, Ganapathy S, Mukherjee S, Srikanthan S, Jolly GP, Anand RS, Prasad SVN (2017) Noncanonical regulation of insulin-mediated ERK activation by phosphoinositide 3-kinase γ. Molecular biology of the cell 28: 3112–3122

Mungai PT, Waypa GB, Jairaman A, Prakriya M, Dokic D, Ball MK, Schumacker PT (2011) Hypoxia triggers AMPK activation through reactive oxygen species-mediated activation of calcium release-activated calcium channels. Mol Cell Biol 31: 3531–3545

Nakamura T, Ranek MJ, Lee DI, Shalkey Hahn V, Kim C, Eaton P, Kass DA (2015) Prevention of PKG1alpha oxidation augments cardioprotection in the stressed heart. J Clin Invest 125: 2468–2472

Nakamura T, Zhu G, Ranek MJ, Kokkonen-Simon K, Zhang M, Kim GE, Tsujita K, Kass DA (2018) Prevention of PKG-1α Oxidation Suppresses Antihypertrophic/Antifibrotic Effects From PDE5 Inhibition but not sGC Stimulation. Circ Heart Fail 11: e004740

Oeing CU, Jun S, Mishra S, Dunkerly-Eyring B, Chen A, Grajeda MI, Tahir U, Gerszten RE, Paolocci N, Ranek MJ et al (2021) MTORC1-Regulated Metabolism Controlled by TSC2 Limits Cardiac Reperfusion Injury. Circ Res

Oeing CU, Nakamura T, Pan S, Mishra S, Dunkerly-Eyring BL, Kokkonen-Simon KM, Lin BL, Chen A, Zhu G, Bedja D et al (2020) PKG1alpha Cysteine-42 Redox State Controls mTORC1 Activation in Pathological Cardiac Hypertrophy. Circ Res 127: 522–533

Pallet N, Legendre C (2013) Adverse events associated with mTOR inhibitors. Expert opinion on drug safety 12: 177–186

Potter CJ, Pedraza LG, Xu T (2002) Akt regulates growth by directly phosphorylating Tsc2. Nat Cell Biol 4: 658–665

Ranek MJ, Kokkonen-Simon KM, Chen A, Dunkerly-Eyring BL, Vera MP, Oeing CU, Patel CH, Nakamura T, Zhu G, Bedja D et al (2019) PKG1-modified TSC2 regulates mTORC1 activity to counter adverse cardiac stress. Nature 566: 264–269

Takimoto E, Champion HC, Li M, Belardi D, Ren S, Rodriguez ER, Bedja D, Gabrielson KL, Wang Y, Kass DA (2005) Chronic inhibition of cyclic GMP phosphodiesterase 5A prevents and reverses cardiac hypertrophy. NatMed 11: 214–222

Takimoto E, Koitabashi N, Hsu S, Ketner EA, Nagayama T, Bedja D, Gabrielson K, Blanton R, Siderovski DP, Mendelsohn ME et al (2009) RGS2 mediates cardiac compensation to pressure-overload and anti-hypertrophic effects of PDE5 inhibition. J Clin Invest 119: 408–420

Tsai EJ, Liu Y, Koitabashi N, Bedja D, Danner T, Jasmin JF, Lisanti MP, Friebe A, Takimoto E, Kass DA (2012) Pressure-overload-induced subcellular relocalization/oxidation of soluble guanylyl cyclase in the heart modulates enzyme stimulation. Circ Res 110: 295–303

Um SH, Frigerio F, Watanabe M, Picard F, Joaquin M, Sticker M, Fumagalli S, Allegrini PR, Kozma SC, Auwerx J et al (2004) Absence of S6K1 protects against age- and diet-induced obesity while enhancing insulin sensitivity. Nature 431: 200–205

Valvezan AJ, Manning BD (2019) Molecular logic of mTORC1 signalling as a metabolic rheostat. Nat Metab 1: 321–333

Wang Y, Viscarra J, Kim SJ, Sul HS (2015) Transcriptional regulation of hepatic lipogenesis. Nature reviews Molecular cell biology 16: 678–689

Yu Q, Gao F, Ma XL (2011) Insulin says NO to cardiovascular disease. Cardiovasc Res 89: 516–524

Zhang HH, Huang J, Duvel K, Boback B, Wu S, Squillace RM, Wu CL, Manning BD (2009a) Insulin stimulates adipogenesis through the Akt-TSC2-mTORC1 pathway. PloS one 4: e6189

Zhang HH, Huang J, Düvel K, Boback B, Wu S, Squillace RM, Wu CL, Manning BD (2009b) Insulin stimulates adipogenesis through the Akt-TSC2-mTORC1 pathway. PloS one 4: e6189

